# A *Solanum lycopersicoides* reference genome facilitates biological discovery in tomato

**DOI:** 10.1101/2020.04.16.039636

**Authors:** Adrian F. Powell, Lance E. Courtney, Maximilian H.-W. Schmidt, Ari Feder, Alexander Vogel, Yimin Xu, David A. Lyon, Kathryn Dumschott, Marcus McHale, Ronan Sulpice, Kan Bao, Asha Duhan, Asis Hallab, Alisandra K. Denton, Lukas A. Mueller, Saleh Alseekh, Jie Lie, Cathie Martin, Alisdair R. Fernie, Sarah R. Hind, Gregory B. Martin, Zhangjun Fei, James J. Giovannoni, Susan R. Strickler, Björn Usadel

**Affiliations:** Boyce Thompson Institute, Ithaca, New York 14853; Institute for Biology I, BioSC, RWTH Aachen University, 52474 Aachen, Germany; IBG-4 Bioinformatics, Forschungszentrum Jülich, 52428 Jülich, Germany; Plant Biology Section, School of Integrative Plant Sciences, Cornell University, Ithaca, NY 14853, U.S.A; Plant Systems Biology Lab, National University of Ireland, Galway, Ireland; Max-Planck-Institut für Molekulare Pflanzenphysiologie, Am Mühlenberg 1, 14476, Potsdam-Golm, Germany; Center of Plant Systems Biology and Biotechnology, 4000 Plovdiv, Bulgaria; Metabolic Biology Department, The John Innes Centre, Norwich Research Park, Norwich, NR4 7UH, UK; Department of Crop Sciences, University of Illinois at Urbana-Champaign, Urbana, IL, 61801, USA; Plant Pathology and Plant-Microbe Biology Section, School of Integrative Plant Science, Cornell University, Ithaca, NY 14853, U.S.A; US Department of Agriculture-Agricultural Research Service, Robert W. Holley Center for Agriculture and Health, Ithaca, NY, USA

## Abstract

Wild relatives of tomato are a valuable source of natural variation in tomato breeding, as many can be hybridized to the cultivated species (*Solanum lycopersicum*). Several, including *Solanum lycopersicoides*, have been crossed to *S. lycopersicum* for the development of ordered introgression lines (ILs). Despite the utility of these wild relatives and their associated ILs, limited finished genomes have been produced to aid genetic and genomic studies. We have generated a chromosome-scale genome assembly for *Solanum lycopersicoides* LA2951 using PacBio sequencing, Illumina, and Hi-C. We identified 37,938 genes based on Illumina and Isoseq and compared gene function to the available cultivated tomato genome resources, in addition to mapping the boundaries of the *S. lycopersicoides* introgressions in a set of cv. VF36 x LA2951 introgression lines (IL). The genome sequence and IL map will support the development of *S. lycopersicoides* as a model for studying fruit nutrient/quality, pathogen resistance, and environmental stress tolerance traits that we have identified in the IL population and are known to exist in *S. lycopersicoides*.

## Introduction

Tomato is the most widely consumed fruit crop with the greatest value world-wide. It is relatively rich in essential nutrients, particularly provitamin A, folate, vitamin C, vitamin E and vitamin K, iron, and calcium although these usually remain below their theoretical maximum levels because they are rarely targets for breeding. In addition, yield is often reduced significantly by losses caused by adverse environmental conditions, disease, pest damage, and post-harvest loss. The narrow germplasm base currently deployed in most breeding programs limits the potential for tomato improvement. Close wild relatives present opportunities to add enormous genetic diversity to tomato breeding programs and the means to identify and study genes that underpin useful novel variation. At least 14 wild relatives can be crossed to cultivated tomato, with varying degrees of difficulty, and have been used for decades in breeding programs (Grandillo *et al.*, 2011). *S. lycopersicoides*, belongs to an outgroup to the tomato clade, and shows enhanced cold tolerance (Zhao *et al.*, 2005), increased anthocyanin content (Rick et al. 1994), exceptional resistance to *Botrytis cinerea* (Guimarães, Chetelat and Stotz, 2004; Davis *et al.*, 2009; Smith *et al.*, 2014) and resistance to *Pseudomonas* (Mazo-Molina *et al.*, 2019). While introgression lines (ILs) are available that can assist identification of alleles and mapping of loci conferring beneficial traits from *S. lycopersicoides* (Canady, Meglic and Chetelat, 2005), few loci have been cloned partly due to a lack of a reference genome assembly for this species. Despite the importance of wild accessions in tomato breeding, only one such species, *S. pennellii*, has a reference quality genome (Bolger *et al.*, 2014; Schmidt *et al.*, 2017). Four species, *S. pimpinellifolium* (Tomato Genome Consortium, 2012; Razali *et al.*, 2017), *S. galapagense* (Strickler *et al.*, 2015), *S. arcanum* and *S. habrochaites* (The 100 Tomato Genome Sequencing Consortium *et al.*, 2014) have draft *de novo* assemblies with small contig size and some have no gene annotation.

Of the biparental tomato IL populations developed to date, *S. lycopersicoides* LA2951 x *S. lycopersicum* cv. VF36 ILs (Canady, Meglic and Chetelat, 2005) represents the widest cross. Due to the lack of a *S. lycopersicoide*s genome, introgression boundaries of these lines are typically defined based on the published tomato reference genome (Tomato Genome Consortium, 2012). Genetic mapping resolution depends on these boundaries being well-defined and assumes lack of genome rearrangements or other modifications. Traditionally boundaries were defined by PCR markers dispersed across the genome, but more recent approaches increase precision by using SNPs derived from resequencing or RNA-Seq data (Gonda *et al.*, 2019) (Chitwood *et al.*, 2013). By basing introgression coordinates on the Heinz reference genome in wide crosses, information regarding differences in genome size is missing, which is relevant when comparing cultivated tomato to wild accessions that often have larger genome sizes. While many differences in genome size are accounted for by repetitive elements, gene gain and loss are also important to consider especially in regard to rapidly evolving genes involved in abiotic stress tolerances and resistance to biotic challenges.

To improve understanding of the genes responsible for agriculturally important traits and increase the utility of wild species introgression lines, we generated a chromosome scale, reference genome and annotation for *S. lycopersicoides* LA2951. We compared the genome to *S. pennellii* and *S. lycopersicum* to identify unique features that may be key to understanding the genetics of abiotic and biotic stress tolerance in *S. lycopersicoides*. Using RNA-seq, we mapped the introgressions of the ILs to both parental genomes to refine the genetic map of the population. The genome and associated map will facilitate improved understanding of the genes responsible for agriculturally important traits enabling their targeted introduction into tomato breeding lines.

## Results

### Genome assembly and feature prediction

*S. lycopersicoides* LA2951 was selected for sequencing due to its traits of interest and the existence of introgression lines previously generated using this accession (Canady, Meglic and Chetelat, 2005). We assembled approximately 17 million PacBio reads with an average length of 6.29 kbp totaling 107 Gbp using two different genome assembly strategies relying on Canu (Koren *et al.*, 2017) and Falcon (Chin *et al.*, 2016). Canu produced an assembly with 17,507 contigs and an N50 value of 139,475. This was the more contiguous assembly which captured a larger proportion of the BUSCO set (Waterhouse *et al.*, 2017)(Supplemental table 1). Thus this assembly was selected for further scaffolding using Dovetail Chicago and Hi-C. The final assembly is 1.2 Gb in length (Table 1) and captures 97% of the BUSCO set (Supplemental figure 1). The genome assembly is larger than the *S. lycopersicum* assembly, which was expected based on kmer analysis (Table 1) and from previous observations (Rick *et al.*, 1986). The final assembly N50 was 93.9 Mb and 90% of the assembly was found in 12 pseudomolecules consistent with most of the assembly being captured by pseudochromosomes. Gaps within scaffolds ranged from 1 to 7,590 bp with a median of 970 bp. Approximately 1% of sites were found to be heterozygous, which is not surprising for a self incompatible species.

**Table 1.** Summary statistics of *S. lycopersicoides* and other tomato genome assemblies

|  | *S. lycopersicoides* | *S. pennellii* v2 | *S. lycopersicum*<br>Heinz v 4.0 |
| --- | --- | --- | --- |
| No. of pseudomolecules | 12 | 12 | 12 |
| longest sequence (Mbp) | 133.5 | 109.3 | 90.9 |
| Contig N50 (bp) | 253,764 | 60,347 | 6,007,830 |
| total length (Mbp) | 1,152 | 926 | 782.5 |
| Total size (bp) of unanchored contigs<br>(% of assembly) | 135,089,793 (10.5) | 63,101,713<br>(6.4%) | 9,643,250 (1.2%) |

Figure 1 shows the twelve pseudochromosomes, along with repeat density and gene feature density. About 68% of the sequence contained in the pseudochromosomes was found to consist of repeats (Figure 2, Supplemental table 2). Genome annotation predicted a total of 37,938 putative genes (34,239 located on pseudomolecules) with a mean length of 4,388 bp (Supplemental table 3). Genes had an average of 5.3 exons and a cds length of 1,232. We were able to identify 96% of the BUSCO set in the protein annotation (Supplemental figure 2).

**Figure 1.**
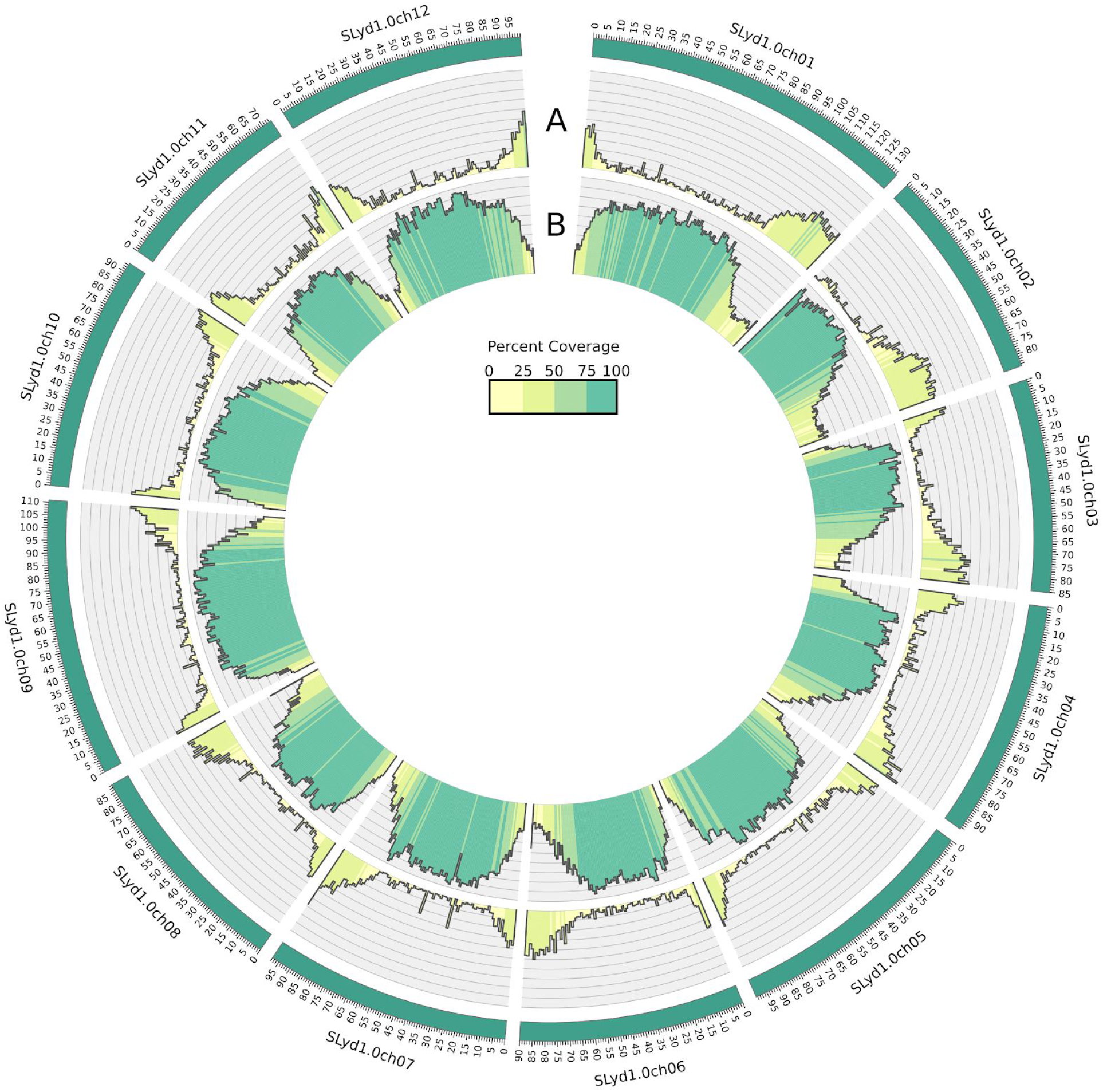
shows the 12 *Solanum lycopersicoides* chromosomes. Circos diagram illustrating the genome of *S. lycopersicoides* with (A) gene densities in the outer data track and (B) repeat densities in the inner data track plotted relative to chromosome position. Densities are calculated for 1 Mb windows and range from 0% to 100%. Different colors are used for values falling within 0% to 25%, 25% to 50%, 50% to 75%, and 75% to 100%.

**Figure 2.**
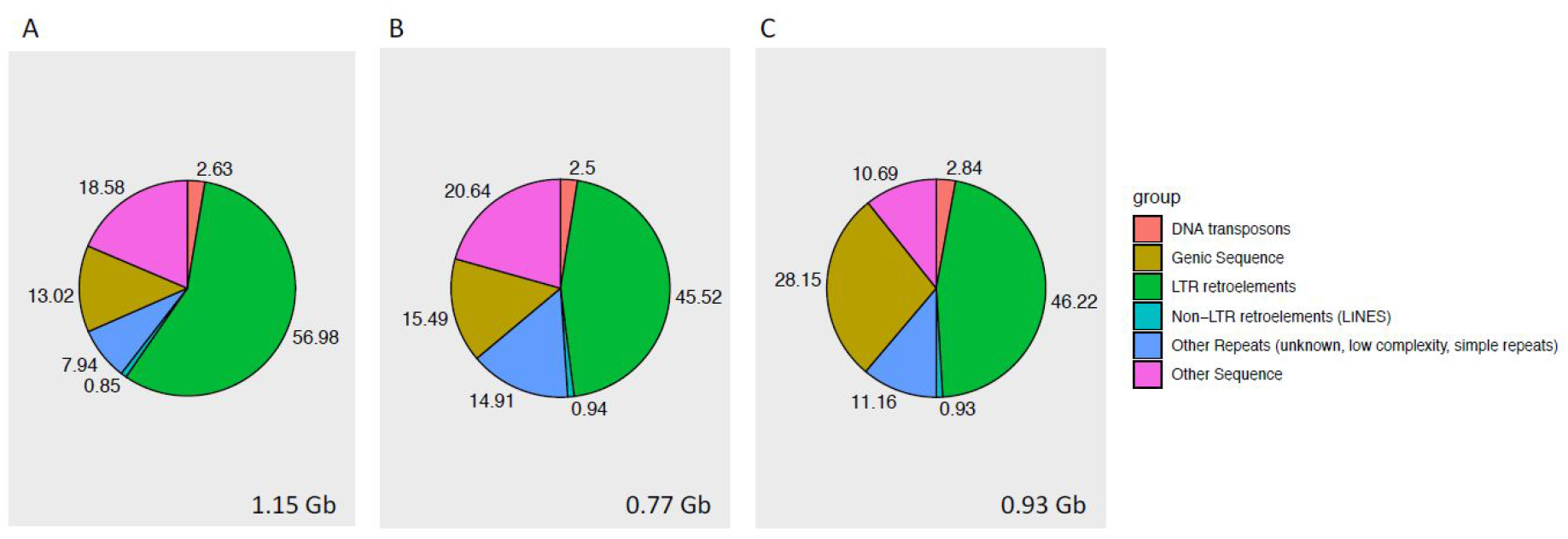
shows the Proportional contributions of component classes to the genomes of (A) *S. lycopersicoides*, (B) *S. lycopersicum*, and (C) *S. pennellii*. These figures relate to the 12 chromosomes for each genome, and the size of each 12-chromosome assembly is shown at the bottom of each panel.

Of all repeat classes, the long terminal repeat retrotransposons (LTR-RT) constituted the greatest proportion (~57%) of the *S. lycopersicoides* genome, as was the case with *S. lycopersicum* and *S. pennellii* genomes (Supplemental tables 2, 4, 5; Figure 2). For LTR-RT elements, *Gypsy*-type LTR-RTs were proportionately more abundant than *Copia*-type elements in each of the three genomes (Supplemental Tables 2, 4, 5). This result had been reported previously for *S. lycopersicum* and *S. pennellii* (Bolger *et al.*, 2014). *S. lycopersicoides* also had a greater proportional abundance of younger LTR-RTs (with insertion times < 1MYA) compared to the other two genome assemblies (Supplemental figure 3).

### Functional annotation and gene families

For annotation the automated functional annotation pipeline Mercator was used which reached classification and annotation rates in line with high quality genomes (Schwacke *et al.*, 2019). The Mercator classification was used to investigate potential gene family expansion in the *Solanum lycopersicoides* genome assembly. This showed that in the case of the histone MLK kinase, there are six genes located on chromosomes 1, 8, 10, 11 and two copies on chromosome 12 both in *S. pennellii* and *S.lycopersicum*. However, in *S. lycopersicoides* there were three copies each in tandem configuration on chromosomes 1 and 8 (Supplemental table 6). In the case of protein elongation, there were multiple copies of genes encoding enzymes involved in synthesizing diphthamide, a modified histidine found only in elongation factor-2 (EEF2) including two DPH1 (Solyd06g065750, Solyd06g065840) and two DPH4 (Solyd12g069970, Solyd12g070080) diphthamide synthesis genes in *S. lycopersicoides*, respectively, but only one each in *S. lycopersicum* and *S. pennellii*. Similarly, there were twice as many deoxyhypusine synthase genes (contig2598g050010, Solyd01g058930, Solyd01g058950, Solyd02g069410). Deoxyhypusine synthase catalyzes the first step in conversion of a lysine residue in eIF5A to the non-standard amino acid, hypusine using spermidine. Similarly, there were twice as many deoxyhypusine hydroxylase proteins (catalysing the second step in the synthesis of hypusine) (Solyd08g069360, Solyd08g069410) presumably generated by tandem duplication, encoded by the *S. lycopersicoides* genome compared to *S. lycopersicum*. This hypusine residue is essential for activity of eIF5A.

Interestingly, within secondary metabolism, both the MVA and MEP pathways providing precursors for isoprenoid biosynthesis were expanded in *S. lycopersicoides*. The domesticated tomato and the *S. pennellii* genomes feature three HMG-CoA Synthases each, while the *S. lycopersicoides* genome features five (Solyd08g053200, Solyd08g072110, Solyd08g072130, Solyd08g072140, Solyd12g068050), due to a tandem expansion on chromosome 8. Furthermore, there might be more HMG-CoA reductase genes in *S. lycopersicoides*, but several were detected on unanchored contigs, only. Similarly, in the MEP pathway, in contrast to the domesticated tomato and *S. pennellii*, which feature one gene each encoding the D-xylulose 5-phosphate transporter, DXR 1-deoxy-D-xylulose 5-phosphate reductase, 4-diphosphocytidyl-2-C-methyl-D-erythritol kinase and 4-hydroxy-3-methylbut-2-enyl diphosphate reductase, *S. lycopersicoides* features two genes in each case.

We detected an expansion of some genes involved in flavonoid synthesis and modification. Whilst dihydroflavonol 4-reductase was triplicated (Solyd02g073780, Solyd02g073830, Solyd02g073850) compared to the domesticated tomato and *S. pennellii*, flavonol-3-O-glycoside-rhamnosyltransferase existed in two copies on chromosomes 3 and 5 in the genome of domesticated tomato, *S. pennellii* showed a tandem duplication of the gene on chromosome 3 and *S. lycopersicoides* showed a (near) tandem duplication on both chromosomes (Solyd03g076070, Solyd03g076130, Solyd05g056030, Solyd05g056040). Finally, we detected a putative expansion of the gene family encoding Phenylalanine Ammonia Lyase (PAL) in *S. lycopersicoides*.

In terms of co-enyzme metabolism, the genome featured several expansions in chlorophyll metabolism.

When investigating potential mechanisms for adaptations to an adverse environment, we noted that CAU1 encoding a histone methylase, which is an epigenetic suppressor of Ca signalling involved in stomatal closure, was duplicated in *S. lycopersicoides*, as well as an expansion of the salt overly sensitive pathway. Here, there was one SOS2 kinase which was duplicated in both wild tomato species compared to domesticated tomato.

In addition, the domesticated tomato as well as the *S. pennellii* genome harbors two SOS1 sodium:protein antiporters whereas there were four in *S. lycopersicoides* due to a tandem triplication on chromosome 4.

Related to immunity, we detected an expansion in *S. lycopersicoides* compared to the domesticated tomato genome and *S. pennellii* of genes encoding members of the G protein family (eight versus five), and AGG1/2 G-gamma component (four versus three in *S.pennellii* or two in the domesticated genome) as well as for the LysM-RLK Lyk5 (two in *S.lycopersicoides* Solyd02g083440, Solyd02g083470). Related specifically to effector-triggered immunity, SGT1 exists in four copies (one being unplaced) but only two in the case of *S. pennellii* and domesticated tomato and there is a dramatic expansion of NLR receptor-encoding genes. Other genes encoding LysM-RLKs, which are typically involved in symbiosis or defense response, depending on species (Buendia *et al.*, 2018), were expanded in *S. lycopersicoides* (Solyd02g078930, Solyd09g070440).

A total of 159,589 out of 208,760 proteins from *Capsicum annum, S. tuberosum*, *S. lycopersicoides*, *S. pennellii*, and *S. lycopersicum* were clustered into 25,763 orthogroups (Figure 3). There were 14,270 orthogroups containing 103,354 proteins that included at least one representative from each of the five species. A total of 29 gene families containing 185 genes were found to be unique to *S. lycopersicoides*. A total of 2,429 orthogroups containing 13,419 genes were unique to *Solanum*.

**Figure 3.**
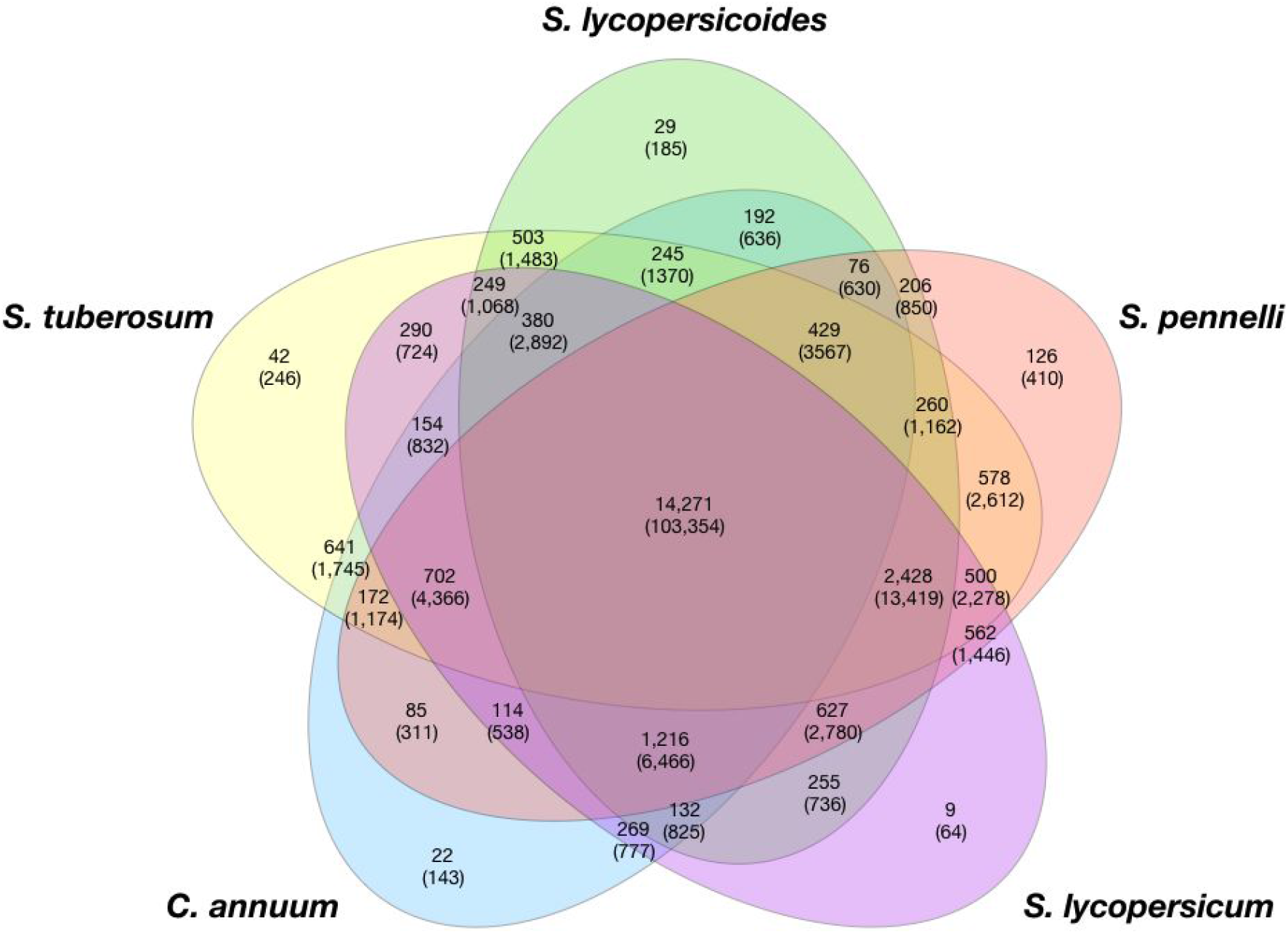
shows number of orthogroups overlapping amongst species. Values in brackets are the number of genes contained in the orthogroups.

### Genome synteny and evolution

Alignment and visualization of paired chromosomal segments of at least 8kb and 92% sequence identity shows the high degrees of synteny between *S. lycopersicoides* and both *S. lycopersicum* and *S. pennellii* (Figure 4). Syntenic dot plots performed at a finer scale identified small inversions relative to the tomato genome (Supplemental figure 4). One inversion found on chromosome 10 is supported by a previously characterized inversion on chromosome 10 in *S. lycopersicoides* relative to *S. lycopersicum* and *S. pennellii* (Canady, Ji and Chetelat, 2006). Another inversion is found near the beginning of chromosome 4.

**Figure 4.**
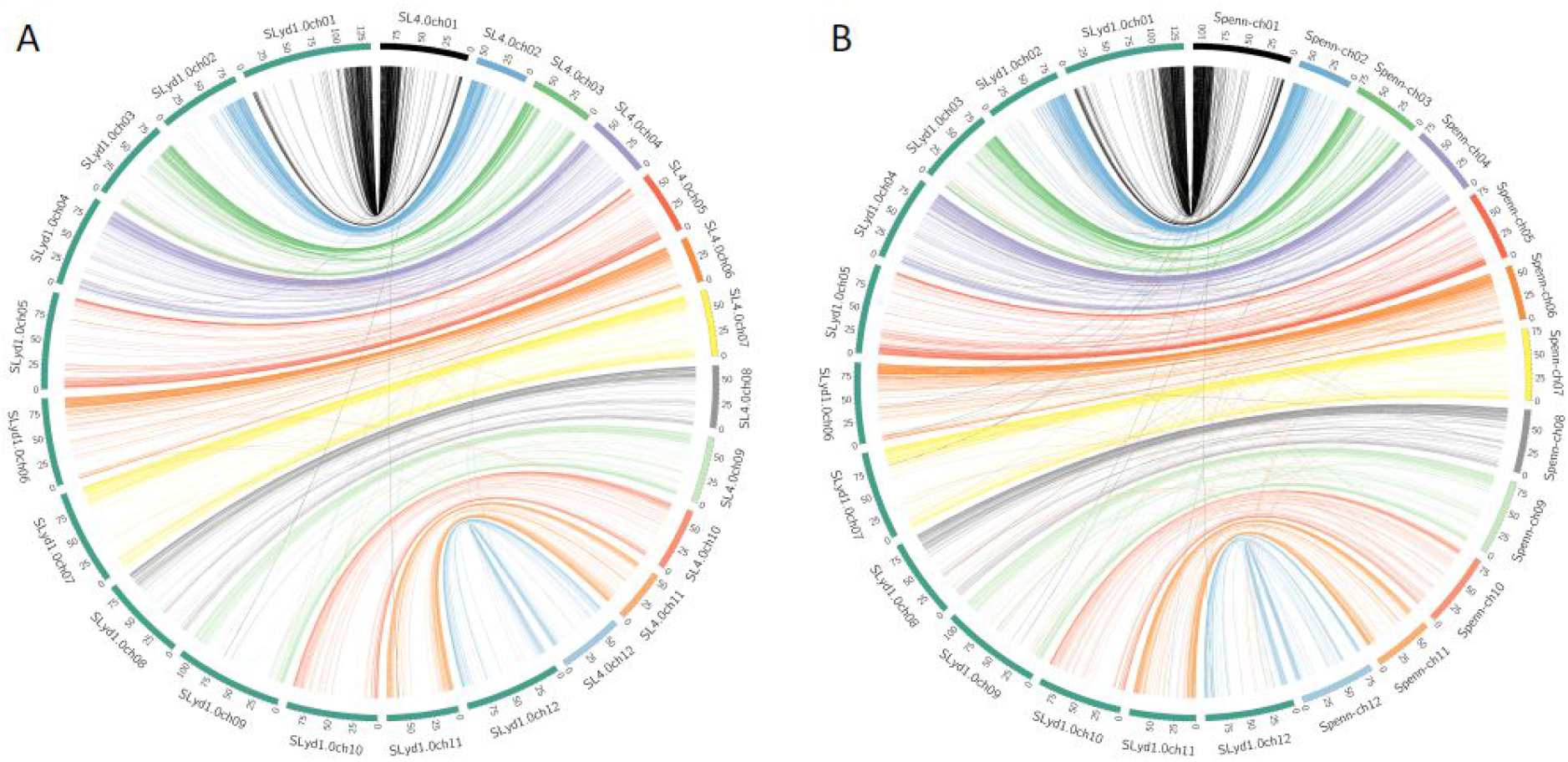
shows syntenic relationships (A) between *S. lycopersicoides* and *S. lycopersicum* as well (B) between *S. lycopersicoides* and *S. pennellii*. Lines in the center of the diagram link aligned sequences from one genome to the other. The color of the lines matches the color given to the corresponding *S. lycopersicum* or *S. pennellii* chromosome in the illustration.

### Introgression mapping

RNA-Seq libraries from 71 unique *S. lycopersicum* cv. VF36 x *S. lycopersicoides* LA2951 IL accessions were used to assemble genotype maps of the population (Sx-Sy). The map based on the current domestic parental reference genome (SL4.0) provides the coordinates delimiting the introgression boundaries within the reference parent background (Sx). An additional map based on the present *S. lycopersicoides* reference provides the coordinates which define the introgressed regions within the wild donor parent (Sy) (Figure 5). Approximately 87% of the *S. lycopersicoides* reference genome chromosomes is represented across these IL accessions. Our analysis revealed numerous previously unidentified introgressed segments and chromosomal features. For example, LA4245 is believed to harbor a single introgression on chromosome 4 (Canady, Meglic and Chetelat, 2005). Our results show that LA4245 harbors an additional ~400kb introgressed region at the distal end of the short arm of chromosome 4. In addition, although this introgression appears on the end of the chromosome within LA4245, it is derived from a region closer to the centromere of chromosome 4 in *S. lycopersicoides* (Sy). This example exposes the risks of low resolution genotype mapping in missing introgressed segments and highlights the potential for error in downstream analyses. Without adequate genotyping, QTL mapping may be suspect. The present genome allowed for high resolution genotype mapping of associated ILs, providing a resource for the community to map QTLs more efficiently and to characterize the variation available from *S. lycopersicoides*.

**Figure 5.**
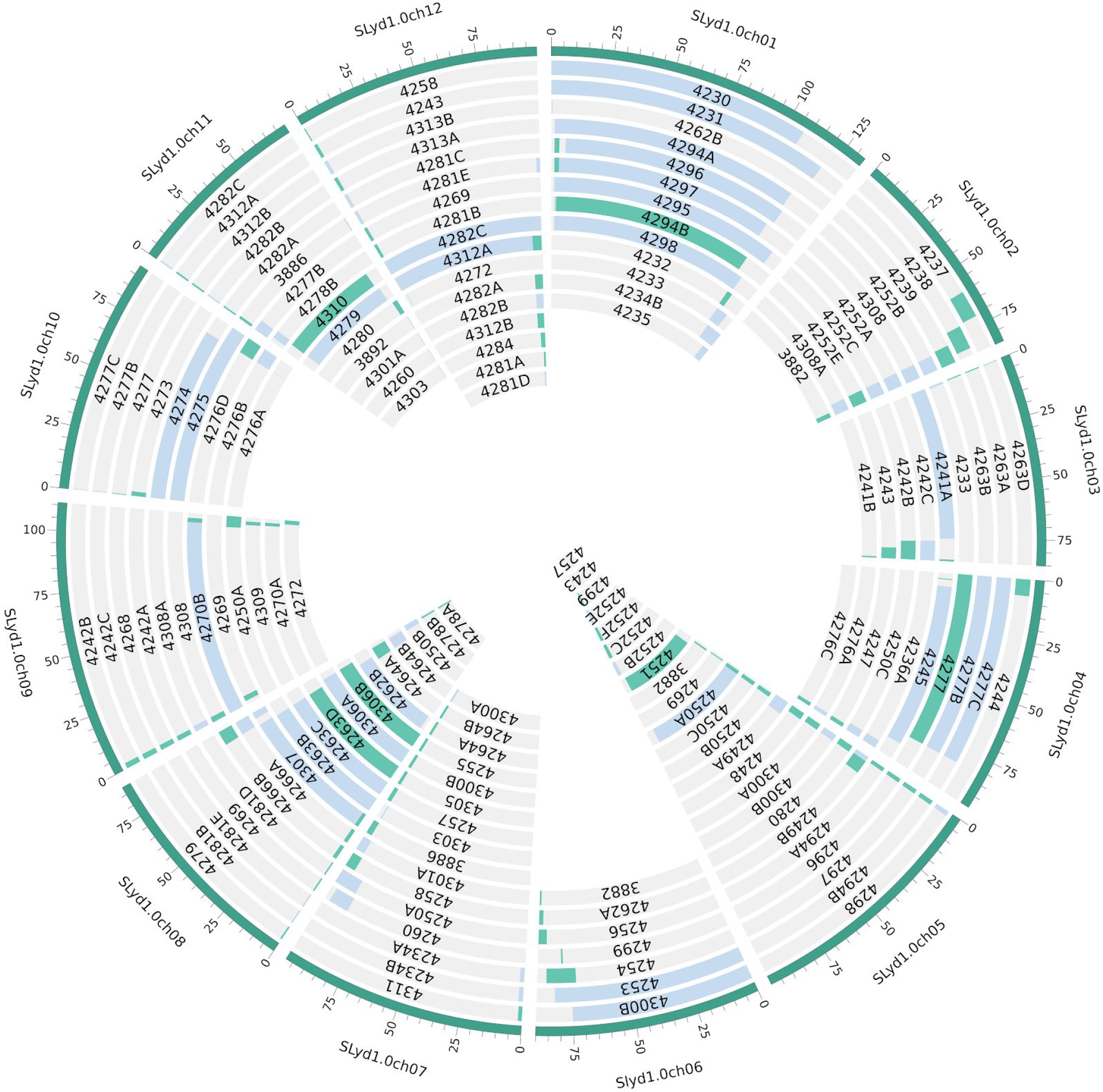
shows the mapping of *S. lycopersicoides* introgression lines onto the genome of *S. lycopersicoides*. Green indicates homozygous *S. lycoperscoides* mapping and blue indicates a mapping of heterozygous regions.

## Discussion

The last few years have seen a tremendous advance in plant genomic sciences driven by technologies such as long read PacBio and lately Oxford Nanopore sequencing (Bolger *et al.*, 2019) as well as new technologies to obtain long range structural information such as optical mapping and Hi-C based information (Schreiber, Stein and Mascher, 2018). Indeed, it has been shown that these technological advances can be used to get (near) chromosome scale assemblies of sub-Gigabase genomes (Belser *et al.*, 2018) and that they are particularly useful to unravel the genomes of wild species which are close relatives of important crops providing information about exotic germplasm and its use (Wu *et al.*, 2018). However, whilst this would be particularly useful in the tomato clade, currently reference-style assemblies exist only for the domesticated tomato (Tomato Genome Consortium, 2012; Razali *et al.*, 2017) and for the wild species *Solanum pennellii* (Bolger *et al.*, 2014; Schmidt *et al.*, 2017). Here, we present a novel high-quality chromosome-scale assembly for the genome of the wild tomato relative *S. lycopersicoides*, which represents an exotic germplasm donor, as together with its sister species *S. sitiens* it makes up the *Solanum* sect. *Lycopersicoides*, which is sister to the sect. *Lycopersicon* clade harboring the domesticated tomato (Knapp and Peralta, 2016). Unlike typical members of sect. *Lycopersicon*, *S. lycopersicoides* does produce large quantities of anthocyanins in the fruit, which might explain it harboring more flavonol-3-O-glycoside-rhamnosyltransferase and flavonol-3-O-glycoside-rhamnosyltransferase genes.

By making use of long read sequencing technology and Dovetail scaffolding, we provide not only a genome, where >90% of the genome can be gathered in 12 chromosome-like scaffolds, but we also show very high completeness of the gene content. This is of particular importance as *S. lycopersicoides* has been used to establish an introgression line population (Canady, Meglic and Chetelat, 2005) which has been used to map several QTL for abiotic and biotic stress tolerance already, given the enhanced resilience of this wild species. Thus, this genome provides a unique reference to fine map and find causative genes underlying QTL when using this IL population. In line with previous marker based data, we also observed an inversion on chromosome 10 of the cultivated tomato relative to the ancestral *S. lycopersicoides* configuration (Pertuzé, Ji and Chetelat, 2002). We found an additional inversion between the cultivated tomato and *S. lycopersicoides* on chromosome 4 which had previously been speculated to be a hotspot as well (Albrecht and Chetelat, 2009).

In addition to improving QTL analyses, we also demonstrate the importance of this genome in assessing gene expression data. Given the large evolutionary distance between *S. lycopersicoides* and the domesticated tomato genome, simply mapping RNA-Seq data to the domesticated gene models showed a low mapping rate. Mapping to closely related relatives has been used (Hekman, Johnson and Kukekova, 2015) and was recommended as a better strategy than transcriptome assemblies (Vijay *et al.*, 2013). Our data shows that this approach can be problematic, especially with distant crosses. While sophisticated bioinformatics pipelines allowing more flexible read alignment and iterative analysis of data and or normalization (Zhou *et al.*, 2019) can reduce these problems, a high-quality reference genome provides the best solution. To improve gene expression analyses within ILs we propose a reference grafting strategy. Using the coordinates from the IL map and both parental reference genomes we assemble synthetic genomes specific for each IL (Supplemental Figure 5). These grafted reference genomes dramatically improve RNA-Seq mapping efficiency within introgressed regions of ILs. IL populations can be powerful tools for both breeding and research communities and the wild reference genome can improve their utility.

Finally, we show that our high-quality genome, together with the available tomato genome data, allows for the assessment of mechanisms of potential adaptation driven by, for example, tandem duplications leading to family expansion.

## Experimental Procedures

### Plant Material

LA2951 seeds were obtained from TGRC and transferred to soil and further cultivated in a greenhouse supplemented with artificial light to a light intensity of at least 200 μmol m^-2^ s^-1^ generated using Phillips hpi-t plus 400w/645 metal-halide lamps for 16 h a day. To preserve the genotype of the used cultivar, one plant was chosen and propagated by cuttings, which were used for DNA extraction.

RNA was prepared from fruit, flowers, and leaves by grinding tissue in liquid N_2_ into very fine powder, aliquoting 200mg of tissue into a 2ml tube, and extracting total RNA using Qiagen RNeasy mini kit. RNA-Seq libraries were prepared following the method described in (Zhong *et al.*, 2011).

### Genome sequencing and assembly

A single *S. lycopersicoides* LA2951 plant was chosen for all genome sequencing efforts. DNA was extracted from young leaves as previously described in (Bolger *et al.*, 2014) and sequenced using PacBio P6C4 technology at Weill Cornell. In addition to this, one Illumina Nextera library, three Illumina TruSeq PCR-based and two Illumina PCR-free libraries were generated and sequenced on an Illumina MiSeq at Research Center Juelich following standard Illumina protocols. The PacBio sequence was assembled using Canu (main version 1.3, commit b147f45b114a9090568d78 fe409557bf5aeeb74f, with the parameter “genomeSize=1.3g”) (Koren *et al.*, 2017) and Falcon (Chin *et al.*, 2016) assemblers (Table 1). BUSCO (Waterhouse *et al.*, 2017) was used to assess the quality of the draft genome assemblies (Figure 2). The Canu genome was polished using Quiver with PacBio reads followed by three iterative rounds of Pilon (Walker *et al.*, 2014) with Illumina paired-end reads and then scaffolded with SSPACE (Boetzer *et al.*, 2011) and Illumina mate pair sequences (Table 1). The assembly was submitted to Dovetail Genomics for further scaffolding using Chicago and Hi-C technologies. Scaffolding gaps were filled with PBJelly (English *et al.*, 2012) using PacBio reads. Heterozygosity was estimated by mapping Illumina reads from genomics DNA to the final assembly with hisat2 and calling SNPs with the GATK (Poplin *et al.*, 2018) pipeline.

### Annotation

For gene model prediction, *de novo* repeats were predicted using RepeatModeler, known protein domains were removed from this set which was then used with RepeatMasker in conjunction with the Repbase library. For gene prediction, RNA was prepared from leaves, flowers, and fruits and sequenced on one lane of Illumina. Reads were mapped to the genome with hisat2 (Kim, Langmead and Salzberg, 2015). Portcullis (Mapleson *et al.*, 2018) and Mikado (Venturini *et al.*, 2018) were used to process the resulting bam files. PacBio IsoSeq data was also generated from a pool of leaves, fruits, and flowers and corrected using the Ice pipeline. Augustus and Snap were trained and implemented through the Maker pipeline, with Iso-Seq, proteins from Swiss-prot, and processed RNA-Seq added as evidence. Functional annotation and classification was performed using the automated Mercator pipeline (Schwacke *et al.*, 2019).

### Repeat Analysis

The genomes of *S. lycopersicoides*, *S. lycopersicum*, and *S. pennellii* were analyzed for LTR retrotransposons using LTRharvest (Ellinghaus, Kurtz and Willhoeft, 2008) with the parameters “-seqids yes-minlenltr 100-maxlenltr 5000-mindistltr 1000-maxdistltr 20000-similar 85 -mintsd 4-overlaps best”. The genomes were then analyzed using LTR_finder (Xu and Wang, 2007) with parameters “-D 20000 -d 1000 -L 5000 -l 100 -p 20 -C -M 0.85 -w 0”. LTR_retriever was used to filter the LTR-RT candidates using default parameters, except that the neutral mutation rate was set at 1.0×10^-8^ using the -u parameter. This neutral mutation rate was selected as it has been used previously for tomatoes (Lin et al., 2014), assuming one generation per year (Beddows *et al.*, 2017).

Candidate miniature inverted-repeat TEs (MITEs) were obtained using MITE-Hunter (Han and Wessler, 2010), with default parameters except for “-P 0.2”. Output candidate MITEs were manually checked for their TSDs and TIRs as suggested in the MITE-Hunter manual. The candidate MITEs were also assigned to superfamilies based on best hits obtained by BLAST against the P-MITE database (http://pmite.hzau.edu.cn/download/), with an e-value cutoff of 1e^-5^. Any candidates that could not be unambiguously classified in this way were classified as unknowns.

The genomes were then masked using the repeat libraries generated by LTR_retriever and MITE-Hunter using Repeatmasker. Additional repeats were then identified *de novo* in the genomes using RepeatModeler. These repeats were classified using blastx against the Uniprot and Dfam libraries and protein-coding sequences were excluded using the script ProtExcluder.pl (Campbell *et al.*, 2014). The masked genomes were then re-masked with Repeatmasker and the corresponding repeat libraries generated by RepeatModeler. Coverage percentages for the various repeat types were calculated using the fam_coverage.pl and fam_summary.pl scripts, both of which are included with LTR_retriever. All coverage percentages were calculated based on the length of the twelve chromosomes as the size for each genome.

### Genome Visualizations and Synteny

Circular genome visualizations were generated using Circos (Krzywinski *et al.*, 2009). For *S. lycopersicoides*, gene and repeat densities were calculated by generating 1MB windows and calculating percent coverage for each feature type (either annotated gene features or repeat elements identified in the repeat analysis) using a bedtools (Quinlan and Hall, 2010; Quinlan, 2014) command of the form: bedtools coverage -a [windows.bed] -b <(sortBed -i [gene or repeat.bed file]) > output. tab.

For synteny analysis, alignment of *S. lycopersicoides* to *S. lycopersicum* (SL4.0) and *S. pennellii* LA0716 v 2 (Bolger *et al.*, 2014) was conducted with nucmer using the parameters --maxgap=500 --mincluster=100, followed by delta-filter -r -q -i 92 -l 8000 and show-coords. Output was then used for links in Circos.

### Gene family analysis

For the Venn diagram, proteins from *C. annuum* v1.55 (Kim *et al.*, 2014), *S. tuberosum* v3.4, *S. lycopersicoides* v 1.0, *S. pennelli* v 2 (Bolger *et al.*, 2014), and *S. lycopersicum* Heinz v 4.0 (Tomato Genome Consortium, 2012) were clustered into orthogroups using OrthoFinder v 2.3.3 (Emms and Kelly, 2015).

### Introgression line maps

RNA-Seq reads were first processed with Trimmomatic-0.36 (Bolger, Lohse and Usadel, 2014) to trim and filter low quality reads. Using Bowtie2 (Langmead and Salzberg, 2012) the reads were then aligned to databases of ribosomal RNA (Quast *et al.*, 2013) and plant virus sequences (Zheng *et al.*, 2017) to filter out non-mRNA reads. The reads were then used to call SNPs following the best practices in the Genome Analysis Toolkit documentation (McKenna *et al.*, 2010). Both *S. lycopersicum* (SL4.0) and *S. lycopersicoides* reference genomes were used to establish a set of SNP markers within the population which distinguish between background and introgressed regions. Using a custom python script, SNPs that did not match either reference genome were filtered out. Only loci that were homozygous for a single parent or heterozygous with one base from each parent were used. This high density genotype marker map was then converted into a map of introgressions using SNPbinner (Gonda *et al.*, 2019). Again, both parental reference genomes were used to map the introgression boundaries within the context of the domesticated background, as well as to positively define the introgressed segments within *S. lycopersicoides*.

## Accession Numbers

The genome sequence is available at Solgenomics.net (Fernandez-Pozo *et al.*, 2015) at https://solgenomics.net/organism/Solanum_lycopersicoides/genome

## Acknowledgements

This work was supported by ERA-CAPS joint call 2 project Regulating Tomato quality through Expression (RegulaTomE) funded under grants DFG US98/7-1 (BU), FE552/29-1 (ARF), and BBSRC BB/N005023/1 (CRM), NSF IOS-1539831 and IOS-1855585 (JJG and ZF), NSF IOS-1546625 (GBM, SRS, and ZF), and the Triad Foundation (SRS, AF, SRH, and GBM). SA and ARF were additionally supported by funding from the Max-Planck-Society and the European Union project PlantaSyst (SGA-CSA No 664621 and No 739582 under FPA No. 664620). BU was supported by BMBF 031A536C. In addition, we want to thank Brian Bell for plant care, Surya Saha and Naama Menda for help with data handling and SGN synchronization and Roger Chetelat and the Tomato Genetics Resource Center for seeds.

## Contributions

BU and SRS managed the project. AFP, MHWS, ZF, JJG, SRS, and BU designed the project. AF, MHS, AV, KD, MM, AD, YX conducted DNA and RNA preparation and sequencing and plant experimental designs. AFP, LEC, MHS, DL, AD contributed new reagents and analytical tools. SA and ARF conducted metabolic analysis and interpretation. AFP, LEC, AKD, ZF, SRS, BU, conducted the data analyses. BU, SRS, AFP, LEC wrote the manuscript with help from all authors.

