## Supplemental Figures and Tables for "A *Solanum lycopersicoides* reference genome facilitates biological discovery in tomato"

Supplemental Figure 1 BUSCO genome analysis

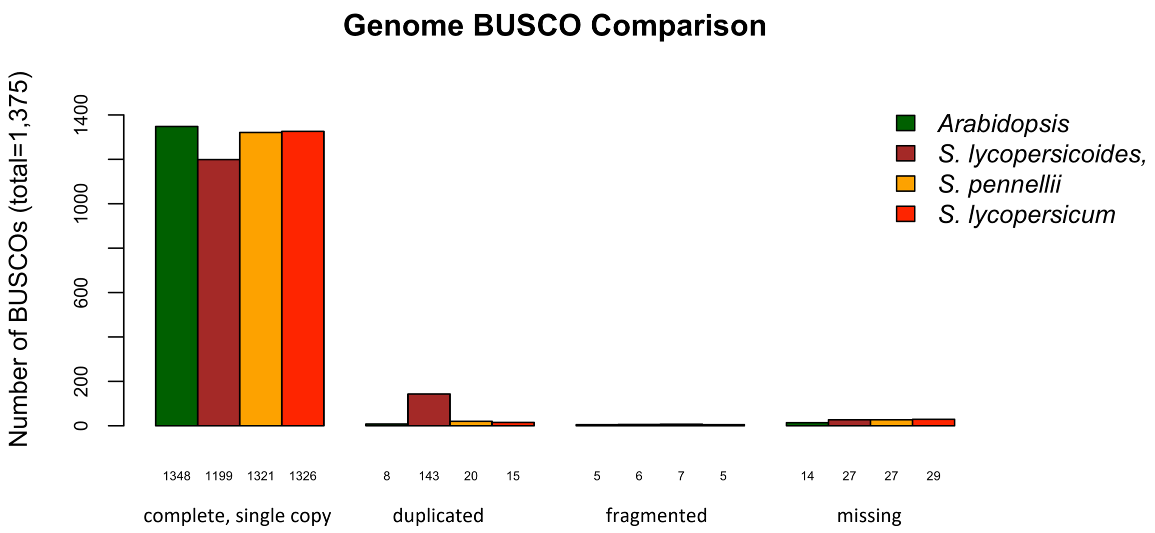


Supplemental Figure 2 BUSCO analysis of annotated protein sets

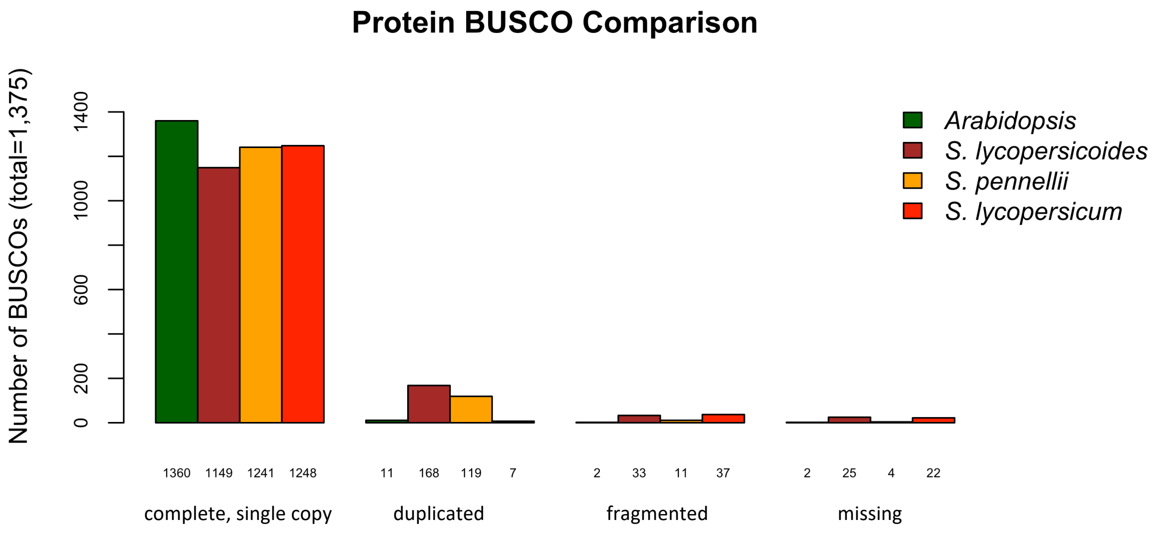

### Supplemental Figure 3 LTR-RT insertion analysis

Age distributions of intact LTR-RTs identified by LTR retriever in the genomes (12 chromosomes) of (A) *S. lycopersicoides*, (B) *S. lycopersicum*, and (C) *S. pennellii*.

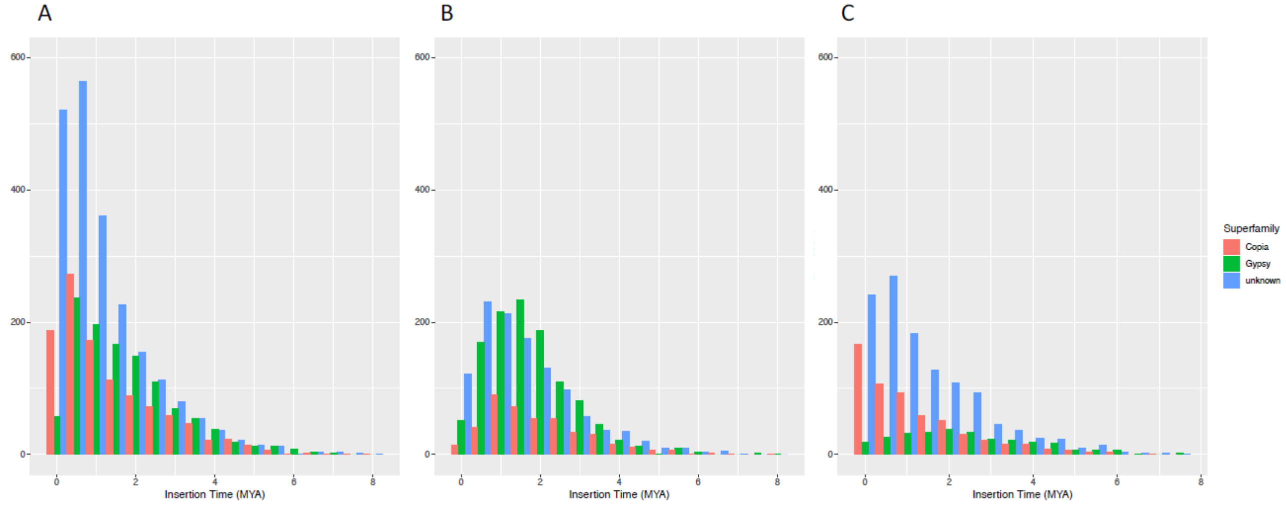


### Supplemental Figure 4 Syntenic Analysis of *S. lycopersicoides* vs *S. lycopersicum*

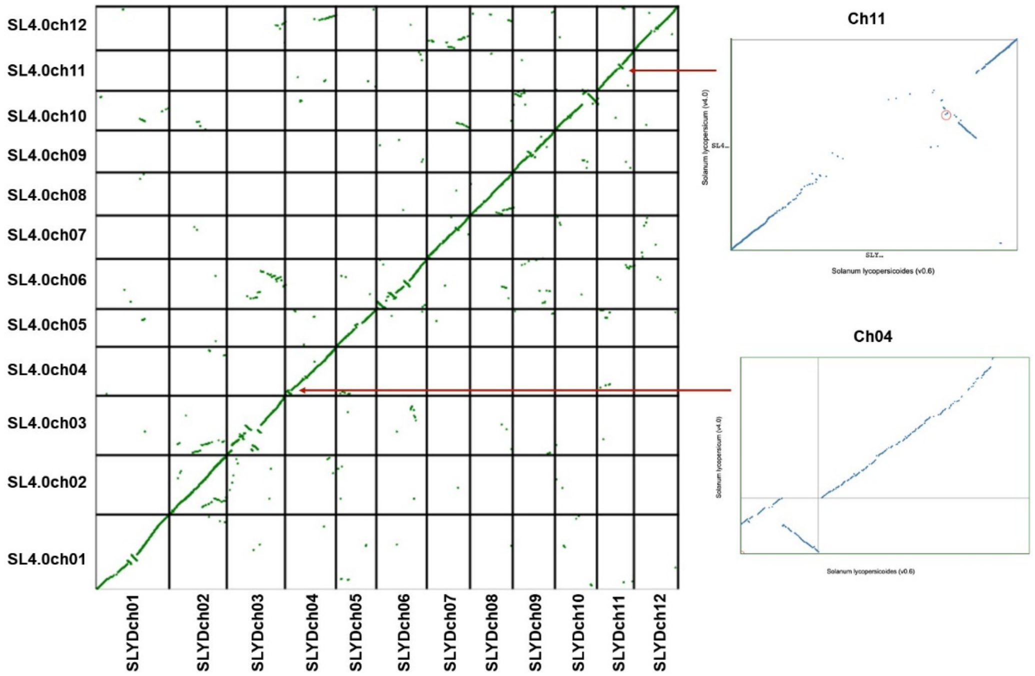

### Supplemental Figure 5 Mapping efficiency of *S. lycopersicoides* IL population

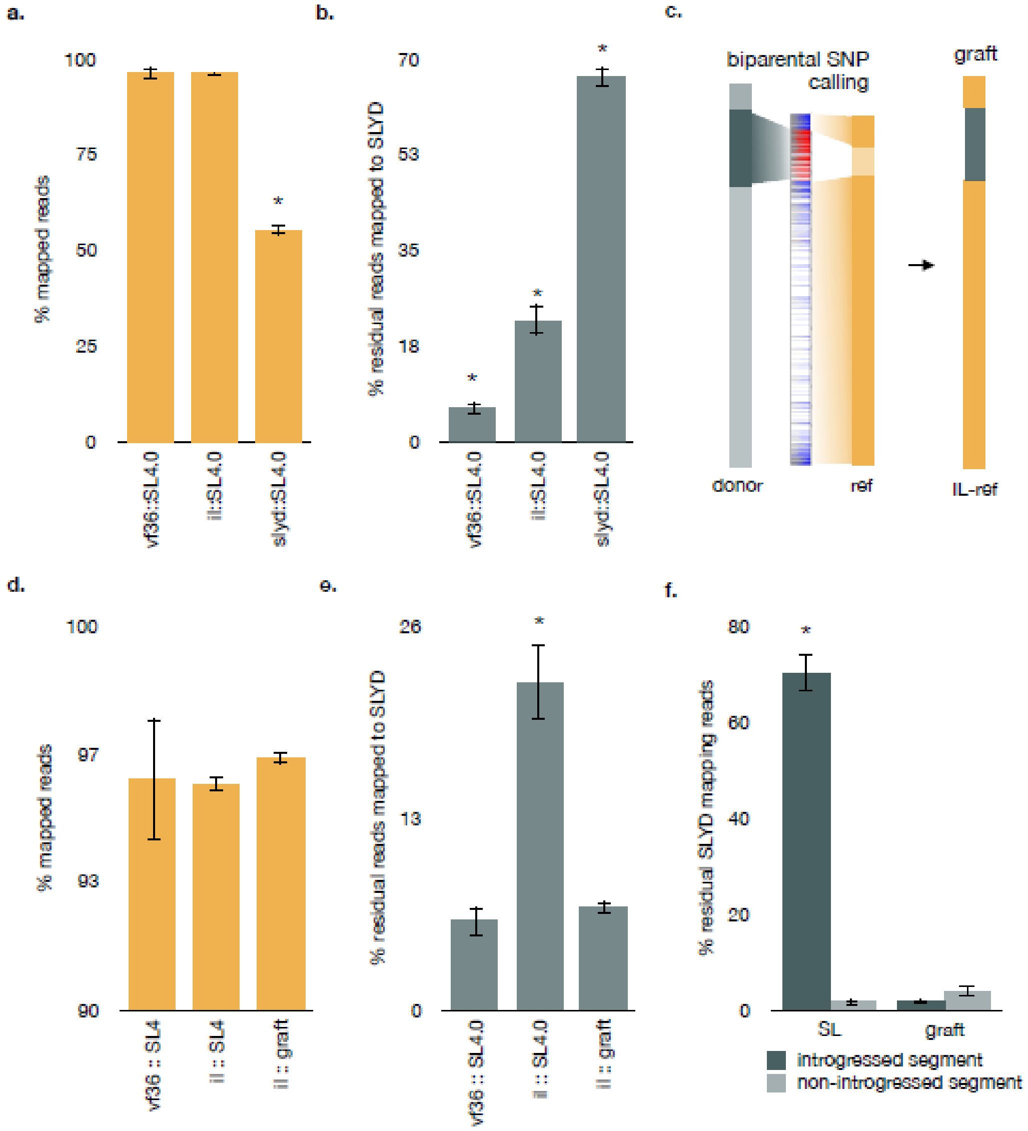


**a,** Overall mapping efficiency of vf36 (domestic reference parent), *S. lycopersicoides* introgression lines (il, n=12), and *S.lycopersicoides* (slyd) RNAseq reads when mapped to the *S. lycopersicum* reference genome SL4.0. **b,** The percent of reads which fail to map to SL4.0, but successfully map to the *S. lycopersicoides* reference genome (SLYD) **c,** Diagram of line specific reference genome grafting using biparental single nucleotide polymorphic (SNP) alignments. **d,** Overall performance of SL4.0 and IL-specific grafted reference assemblies in mapping il reads. **e,** Percent of reads which fail to map to either SL4.0 or the IL-graft, but successfully map to the SLYD reference **f,** Percent of residual (non-SL4.0 or non-IL-graft mapping) reads which map to introgressed segments of SLYD. Error bars represent the standard error of each set: n=6 for vf36 reads, n=4 for slyd reads, n=12 for introgression lines (each line represents an average of at least 3 replicate libraries). One-way ANOVA with Tukey's HSD correction for multiple comparisons was used to determine significance ($p \leq 0.05$).

Supplemental Table 1 Primary assembly statistics

| | Falcon | Canu |
|---|---|---|
| **Total length (Mbp)** | 1,031 | 1,255 |
| **No. of sequences** | 16,350 | 17,507 |
| **Longest sequence (Kbp)** | 2,477 | 3,446 |
| **N50 (bp)** | 130,826 | 139,475 |
| **BUSCO (complete/missing)** | 1,139/246 | 1203/48 |

Supplemental Table 2 Repeat analysis data for the *S. lycopersicoides* (Slyd1.0) genome. Counts and percentages are calculated using the 12 chromosomes in the assembly.

| Classification | Order | Superfamily | No. of TEs | Coverage (bp) | Fraction of genome (%) |
|---|---|---|---|---|---|
| **Class I** | LTR | *Copia* | 92249 | 51271188 | 4.45 |
| | LTR | *Gypsy* | 437886 | 399097526 | 34.64 |
| | LTR | Unknown/Other | 325754 | 206098669 | 17.89 |
| | LINE | *L1* & Other | 30408 | 9818925 | 0.85 |
| **Class II** | TIR | *CACTA* | 4511 | 811613 | 0.07 |
| | TIR | *CMC-EnSpm* | 4534 | 2482189 | 0.22 |
| | TIR | *hAT* | 24007 | 5610654 | 0.49 |
| | Helitron | *Helitron* | 638 | 297722 | 0.03 |
| | TIR | *Mutator* | 44623 | 12469235 | 1.08 |
| | TIR | *PIF-Harbinger* | 10856 | 3177671 | 0.28 |
| | TIR | *Tc1-Mariner* | 27093 | 5242664 | 0.46 |
| | TIR | Unknown | 919 | 216748 | 0.02 |
| **Other - Simple Repeats** | | | 3863 | 1301116 | 0.11 |
| **Other - Unknown Repeats** | | | 410552 | 90194695 | 7.83 |
| **Total** | | | | | 68.40 |

Supplemental Table 3 Annotation metrics

| | *S. lycopersicoides* v1.0 | *S. pennellii* v2 | *S. lycopersicum* v4.0 |
|---|---|---|---|
| **no. of gene models** | 37,938 | 48,923 | 34,075 |
| **Average gene model length (bp)** | 4,388 | 5,962 | 3,571 |
| **Average CDS length (bp)** | 1,232 | 1,558 | 1,027 |
| **Average exons/gene** | 5.3 | 6 | 4.5 |

Supplemental Table 4 Repeat analysis data for the *S. lycopersicum* (Slyc4.0) genome. Counts and percentages are calculated using the 12 chromosomes in the assembly.

| Classification | Order | Superfamily | No. of TEs | Coverage (bp) | Fraction of genome (%) |
|---|---|---|---|---|---|
| **Class I** | LTR | *Copia* | 89814 | 49354289 | 6.39 |
| | LTR | *Gypsy* | 284587 | 220025190 | 28.47 |
| | LTR | Unknown/Other | 134678 | 82411485 | 10.66 |
| | LINE | *L1* & Other | 22764 | 7250148 | 0.94 |
| **Class II** | TIR | *CACTA* | 1968 | 323508 | 0.04 |
| | TIR | *CMC-EnSpm* | 3830 | 2301067 | 0.30 |
| | TIR | *hAT* | 11046 | 2639124 | 0.34 |
| | Helitron | *Helitron* | 539 | 294078 | 0.04 |
| | TIR | *Mutator* | 31042 | 10845303 | 1.40 |
| | TIR | *PIF-Harbinger* | 3701 | 880013 | 0.11 |
| | TIR | *Tc1-Mariner* | 9850 | 1750009 | 0.23 |
| | TIR | Unknown | 519 | 313458 | 0.04 |
| **Other - Simple Repeats** | | | 2065 | 431326 | 0.06 |
| **Other - Unknown Repeats** | | | 421337 | 114826198 | 14.86 |
| **Total** | | | | | 63.87 |

Supplemental Table 5 Repeat analysis data for the *S. pennellii* genome (version from Bolger et al., 2014; available at https://solgenomics.net/organism/Solanum_pennellii/genome). Counts and percentages are calculated using the 12 chromosomes in the assembly.

| Classification | Order | Superfamily | No. of TEs | Coverage (bp) | Fraction of genome (%) |
|---|---|---|---|---|---|
| **Class I** | LTR | *Copia* | 97480 | 41738090 | 4.51 |
| | LTR | *Gypsy* | 414809 | 288196376 | 31.11 |
| | LTR | Unknown/Other | 221842 | 98256230 | 10.61 |
| | LINE | *L1* & Other | 27829 | 8630199 | 0.93 |
| **Class II** | TIR | *CACTA* | 4985 | 940926 | 0.10 |
| | TIR | *CMC-EnSpm* | 5190 | 2794412 | 0.30 |
| | TIR | *hAT* | 16803 | 4397608 | 0.47 |
| | Helitron | *Helitron* | 739 | 420122 | 0.05 |
| | TIR | *Mutator* | 43058 | 12107956 | 1.31 |
| | TIR | *PIF-Harbinger* | 8539 | 2033138 | 0.22 |
| | TIR | *Tc1-Mariner* | 17938 | 3528403 | 0.38 |
| | TIR | Unknown | 564 | 101475 | 0.01 |
| **Other - Simple Repeats** | | | 5513 | 2126207 | 0.23 |
| **Other - Unknown Repeats** | | | 388638 | 101292712 | 10.93 |
| **Total** | | | | | 61.16 |
